# CNOT6 deadenylase safeguards postnatal growth and metabolic transition via *Fgf21* mRNA decay

**DOI:** 10.64898/2026.02.17.706459

**Authors:** Moawiz Saeed, Mengwei Zang, David Gius, Sakie Katsumura, Masahiro Morita

## Abstract

Postnatal growth and development require precise coordination of growth and metabolism to meet the biosynthetic and energetic demands of rapidly expanding organs. Fibroblast growth factor 21 (FGF21) serves as a key endocrine regulator linking nutrient availability to systemic growth control in early life and metabolic homeostasis in adulthood. Here, we identify the CCR4-NOT deadenylase complex subunit CNOT6, but not its paralog CNOT6L, as an essential post-transcriptional regulator of neonatal growth and metabolism. Loss of *Cnot6* results in severe growth retardation, multi-organ hypoplasia, and increased perinatal mortality. Surviving *Cnot6* knockout mice display markedly reduced body and organ size that gradually normalizes by adulthood, indicating developmental compensation. Mechanistically, *Cnot6* deficiency elevates hepatic *Fgf21* mRNA expression, suppresses the IGF1-IGFBP1 axis, and reprograms liver transcriptional networks controlling lipid and glucose metabolism and apoptosis. These changes are accompanied by increased ketone body production, suggesting enhanced fatty acid oxidation. Together, our findings uncover a previously unrecognized role of CNOT6 in limiting FGF21 expression to preserve anabolic metabolism during the neonatal period. This work establishes the CNOT6-FGF21 axis as a molecular checkpoint that couples mRNA decay with hormonal and metabolic coordination required for healthy postnatal growth.

## INTRODUCTION

Fetal and postnatal growth and development are tightly coordinated processes that integrate cellular proliferation, differentiation, and metabolism to ensure tissue expansion and organ maturation (Black et al., 2013; Prendergast and Humphrey, 2014; Victora et al., 2021). The transition from fetal to postnatal and later developmental stages requires systemic metabolic reprogramming of anabolic pathways and energy utilization, driving accelerated growth and metabolic remodeling of multiple organs, including the liver, muscle, adipose tissue, and brain. Disruptions in metabolic adaptation during these stages limit the synthesis of key macromolecules, such as proteins, lipids, nucleic acids, and ATP, leading to impaired growth and organ development with long-lasting consequences, even if catch-up growth occurs later. Epidemiological studies link restricted infant growth to a higher lifelong risk of metabolic disorders, such as obesity and type 2 diabetes (Dewey and Begum, 2011). Although nutritional interventions have reduced the number of stunted infants, the overall prevalence of stunting remains high (Bhutta et al., 2013; Dewey et al., 2021), reflecting the complex interplay of nutritional, hormonal, and metabolic mechanisms. These findings highlight the need to elucidate the molecular mechanisms that coordinate metabolism and growth during the fetal and postnatal stages.

Several key regulators have been implicated in fetal and postnatal growth, including endocrine factors such as growth hormone (GH), insulin-like growth factor 1 (IGF1), and fibroblast growth factor 21 (FGF21), as well as nutrient-sensing pathways such as mTOR and AMPK (Flippo and Potthoff, 2021; Gonzalez et al., 2020; Hage et al., 2021; Liu and Sabatini, 2020; Ranke and Wit, 2018). Among these regulators, FGF21 has emerged as a pivotal endocrine mediator linking metabolism to postnatal growth (Flippo and Potthoff, 2021; Inagaki et al., 2008; Kliewer and Mangelsdorf, 2019). FGF21 is primarily secreted by the liver and orchestrates systemic metabolic adaptation by regulating lipid and glucose metabolism, insulin sensitivity, and energy expenditure in response to nutrient availability (Flippo and Potthoff, 2021; Kliewer and Mangelsdorf, 2019). During postnatal development in humans, elevated FGF21 expression is associated with reduced body size (Fazeli et al., 2010; Guasti et al., 2014; Lee et al., 2023; Mericq et al., 2014; Mistry et al., 2023). In mice, FGF21 overexpression causes significant postnatal growth inhibition by inducing GH resistance (Inagaki et al., 2008; Zhang et al., 2012). Mechanistically, FGF21 attenuates the IGF1-IGFBP1 axis by suppressing hepatic IGF1 expression while upregulating IGFBP1 expression, collectively impairing linear growth in early life. In contrast, FGF21 exerts beneficial metabolic effects in adults, and multiple FGF21 analogs are currently being tested in clinical trials for MASH and hypertriglyceridemia (Chui et al., 2024; Geng et al., 2020). These findings position FGF21 as a critical regulator at the intersection of nutrient metabolism and developmental growth control.

Recent studies have identified the CCR4-NOT complex as a key regulator of hepatic FGF21 expression and broader metabolic gene networks (Katsumura et al., 2022; Morita et al., 2019). The CCR4-NOT complex is the major eukaryotic deadenylase machinery responsible for shortening the 3’ poly(A) tail of mRNAs, thereby promoting the decay of specific mRNAs and fine-tuning gene expression (Collart et al., 2024; Passmore and Coller, 2022; Shirai et al., 2014). This multi-subunit complex comprises the catalytic deadenylases CNOT6, CNOT6L, CNOT7, and CNOT8; the scaffold protein CNOT1; and additional regulatory subunits, including CNOT2, CNOT3, CNOT9, CNOT10, and CNOT11. Among these, either CNOT6 or CNOT6L serves as the primary deadenylase in mammals and acts as a rate-limiting enzyme for mRNA turnover (Collart et al., 2024; Katsumura et al., 2022; Morita et al., 2007; Passmore and Coller, 2022). In adult mice, CNOT6L has been shown to regulate systemic metabolism, including energy expenditure and lipid metabolism by promoting the degradation of Fgf21 mRNA (Katsumura et al., 2022; Morita et al., 2019). Despite these findings, the physiological role of the CCR4-NOT complex in postnatal growth and development and its link to key metabolic regulators such as FGF21 remain largely unknown.

Here, we demonstrate that CNOT6 deadenylase, but not its paralog CNOT6L, plays a critical role in the post-transcriptional machinery securing early-life growth and metabolism. Using a *Cnot6* knockout (KO) mouse model, transcriptomic profiling, and metabolic blood analyses, we demonstrate that *Cnot6* KO mice show markedly reduced postnatal body size, impaired organ growth, and elevated neonatal mortality. These defects are accompanied by increased hepatic FGF21 expression, suppression of the IGF1-IGFBP1 axis, and enhanced fat catabolism. Collectively, our findings uncover an unrecognized role for the CNOT6-FGF21 axis in orchestrating hormonal and metabolic pathways essential for postnatal growth, providing new insights into the molecular basis of early-life growth failure and metabolic dysregulation.

## RESULTS

### CNOT6L deletion has no impact on early postnatal growth or organ size

CNOT6L regulates systemic metabolism in adult mice by modulating FGF21 expression (Katsumura et al., 2022), prompting us to hypothesize that CNOT6L also contributes to postnatal growth and metabolism via a similar mechanism. To evaluate the contribution of CNOT6L to postnatal body growth, we monitored body size and weight in *Cnot6l* wild-type (WT) and knockout (KO) mice (Morita et al., 2019) during postnatal development. At the weaning stage (2-3 weeks of age), no apparent differences in overall appearance were observed between male or female *Cnot6l* WT and KO mice (Figures 1A-B). Consistent with this observation, both body weight and nose-to-anus length did not significantly differ between WT and KO mice (Figures 1C-F). Tibia length, an additional indicator of body size, showed no significant difference between WT and KO (Figures 1G-H). Body weight was measured weekly from 3 to 7 weeks of age, and both WT and KO mice showed comparable growth curves (Figures 1I-J). We next assessed organ weights at 2 weeks of age and found no significant differences in the liver, spleen, kidney, heart, lung, or brain between WT and KO mice (Figures S1A-L). Collectively, these results indicate that CNOT6L deletion does not affect body size or organ growth during early postnatal development, suggesting that its physiological role is more critical for metabolic regulation in adulthood.

**Figure 1.**
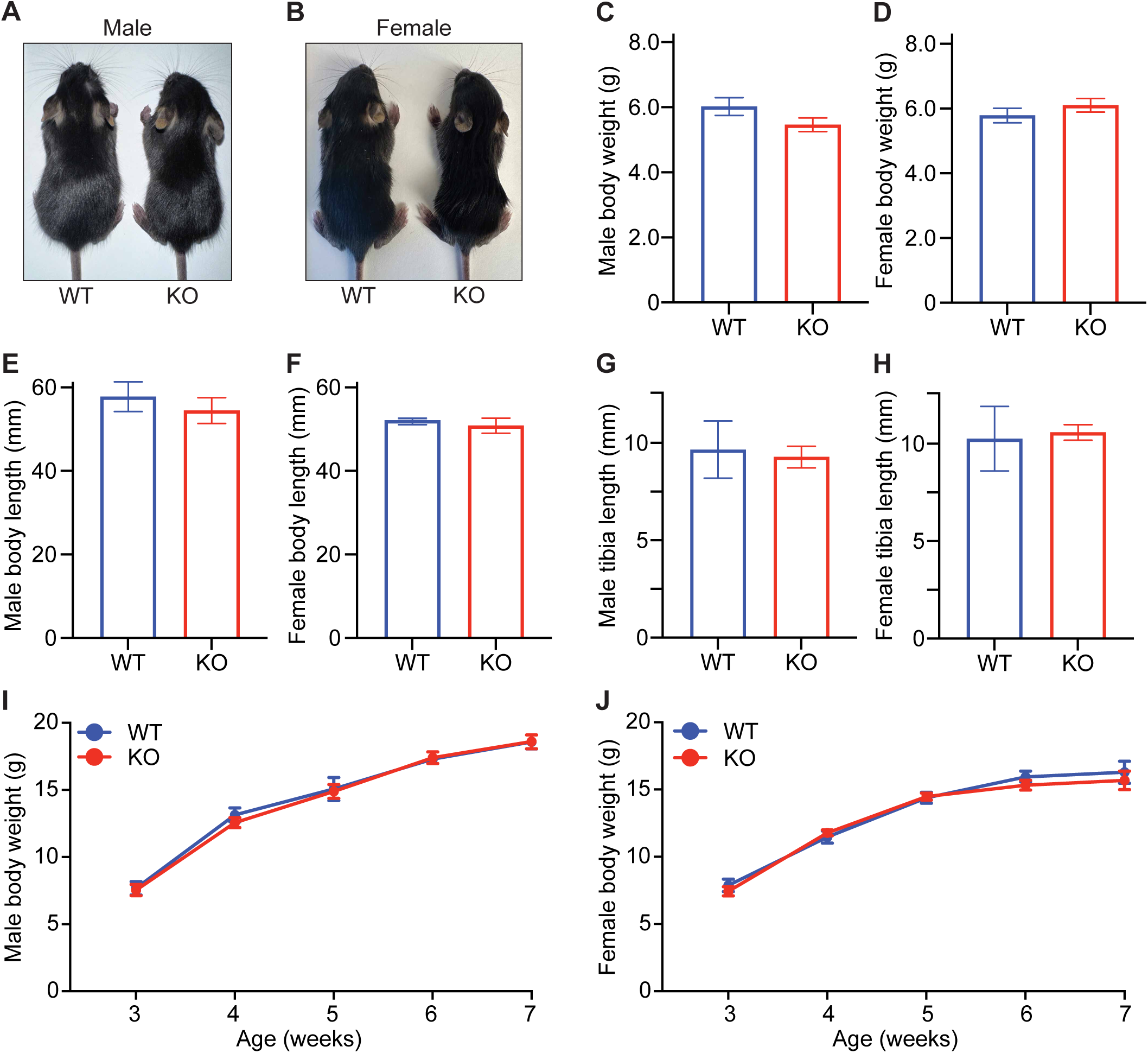
CNOT6L deficiency does not affect postnatal growth or organ size. (A-B) Representative images of 2-week-old male (A) and female (B) *Cnot6l* WT and KO mice. (C-H) Body weights (C-D), body (nose-to-anus) lengths (E-F), and tibia lengths (G-H) of 2-week-old male and female *Cnot6l* WT and KO mice. n = 4-15 per group. (I-J) Growth curves showing body weights of male (I) and female (J) *Cnot6l* WT and KO mice from 3 to 7 weeks of age. n = 3-14 per group. All data are presented as mean ± SEM. See also Figure S1.

### CNOT6 deletion impairs early postnatal survival and leads to growth retardation

CNOT6L deficiency had no effect on early postnatal growth or organ size. Given the close homology between CNOT6L and its paralog CNOT6 (80% sequence identity) (Morita et al., 2007), both of which possess comparable deadenylase activity (Katsumura et al., 2022; Wang et al., 2010), we next examined whether CNOT6 functions as the key CCR4-NOT component governing early postnatal growth and metabolism. To investigate the role of CNOT6 in early development, we generated *Cnot6* KO mice using a targeted gene-trap strategy (Skarnes et al., 2011). In this model, a lacZ operon, a neomycin resistance cassette, and a polyadenylation sequence were inserted between exons 3 and 4 of the *Cnot6* gene (Figure S2A), disrupting normal transcription and resulting in a complete loss of detectable CNOT6 protein expression (Figures S2B-C). Importantly, male and female *Cnot6* KO mice exhibited dramatic growth impairment. At 2 weeks of age, both male and female Cnot6 KO pups appeared smaller than their WT littermates (Figures 2A-B). Quantitative analysis confirmed significant reductions in body weight, nose-to-anus length, and tibia length in *Cnot6* KO mice of both sexes (Figures 2C-H). Despite these early growth defects, body weight monitoring from 3 to 8 weeks of age revealed that *Cnot6* KO mice gradually caught up to WT levels, with no significant difference by 8 weeks (p = 0.065; Figures 2I-J).

**Figure 2.**
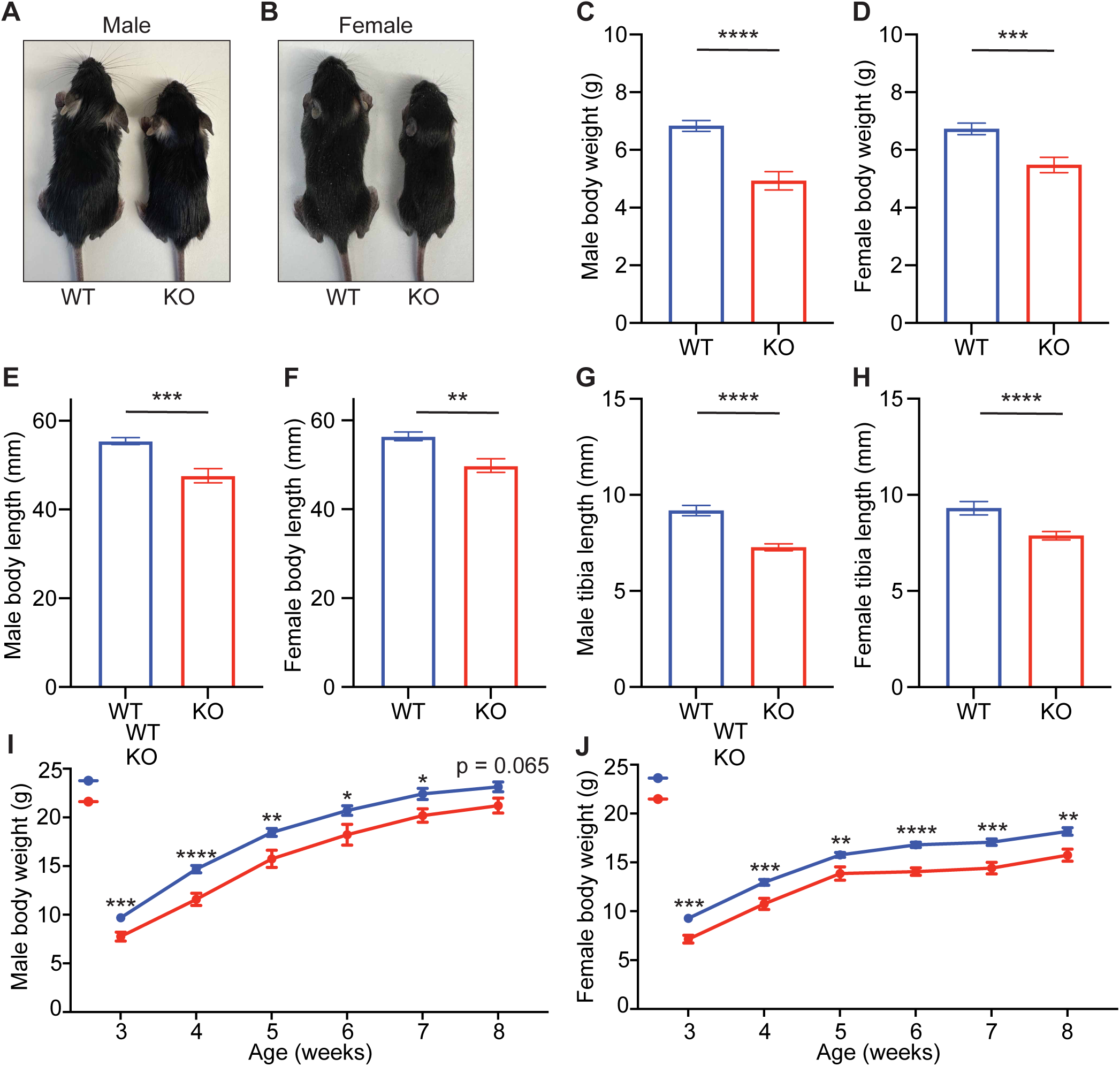
CNOT6 deficiency causes severe growth retardation and postnatal lethality. (A-B) Representative images of 2-week-old male (A) and female (B) *Cnot6* WT and KO mice. (C-H) Body weights (C-D), body (nose-to-anus) lengths (E-F), and tibia lengths (G-H) of 2-week-old male and female *Cnot6* WT and KO mice. n = 6-40 per group. (I-J) Growth curves showing body weights of male (I) and female (J) *Cnot6* WT and KO mice from 3 to 8 weeks of age. n = 3-6 per group. All data are presented as mean ± SEM. *p < 0.05, **p < 0.01, ***p < 0.001, ****p < 0.0001; p values by the Student’s t test (C-H) and one-way ANOVA (I-J). See also Figure S2.

Given the significant reduction in body size observed in *Cnot6* KO mice, we next evaluated the impact of *Cnot6* deletion on organ development. At 2 weeks of age, *Cnot6* KO mice exhibited visibly smaller organs compared with WT littermates (Figures 3A-P). Quantitative analysis revealed significant reductions in the weights of the liver, spleen, kidneys, and heart (Figures 3A-P). These results indicate that CNOT6 is required for proper postnatal organ growth, particularly in highly metabolic organs such as the liver and kidneys.

**Figure 3.**
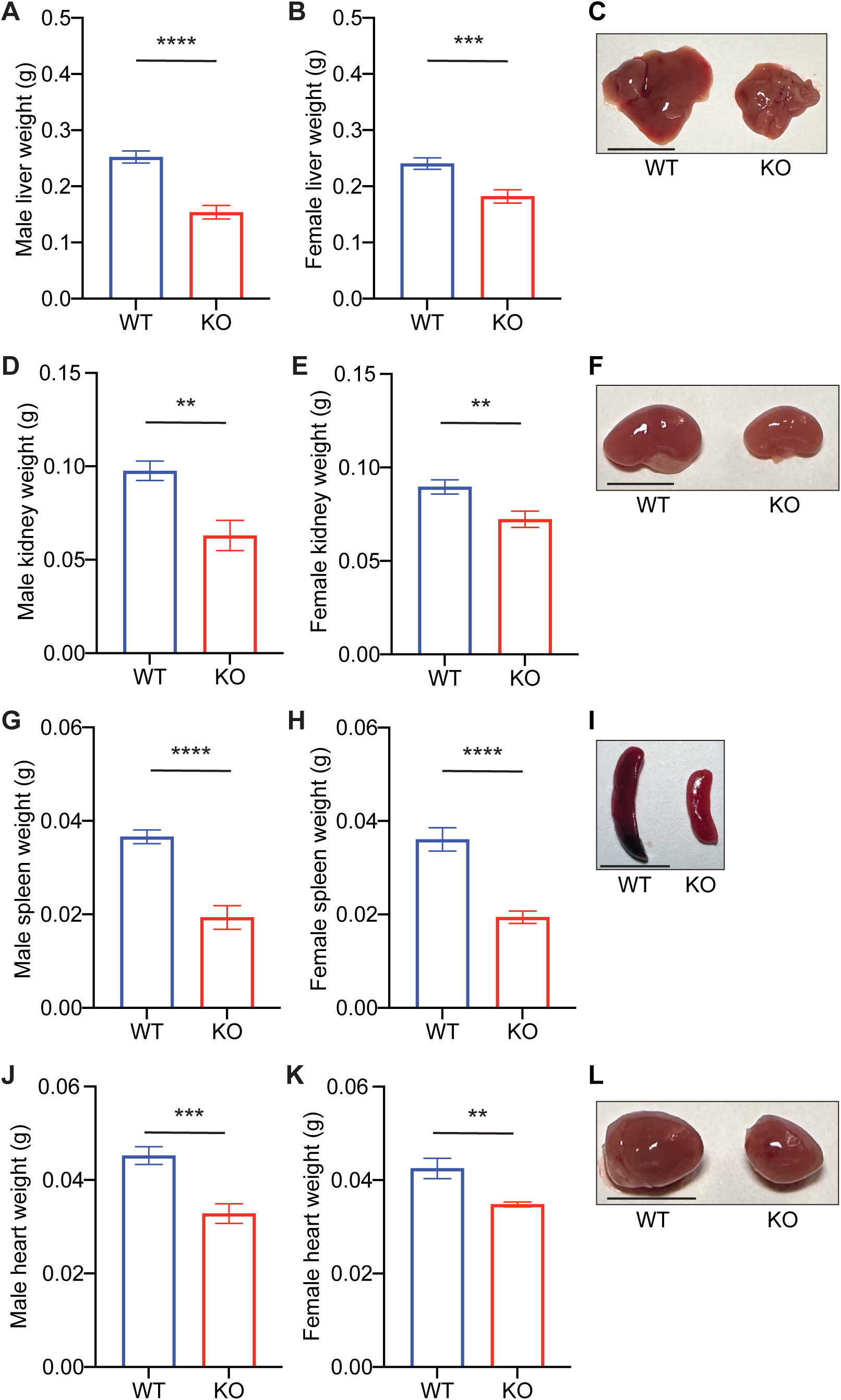
*Cnot6* deletion impairs organ growth during postnatal development. (A-L) Weights and representative images of tissues, including livers (A-C), kidneys (D-F), spleens (G-I), and hearts (J-L) of *Cnot6* WT and KO male and female mice. n = 5-38 per group. Red reference line indicates 10 mm. All data are presented as mean ± SEM. *p < 0.05, **p < 0.01, ***p < 0.001, ****p < 0.0001; p values by the Student’s t test. See also Figure S3

To further assess the growth defects of *Cnot6* deletion on early postnatal survival, we genotyped offspring from *Cnot6* heterozygous (HET) intercrosses at 2 weeks of age. The observed genotype distribution did not follow the expected Mendelian ratio (1 WT: 2 HET: 1 KO) (Figure S3). Specifically, while 113 WT mice were identified, only 37 homozygous KO mice were obtained, corresponding to 32.7% of the expected KO frequency (Figure S3). This finding suggests that a substantial proportion of *Cnot6* KO mice die before 14 days of age. To define the timing of lethality, we genotyped pups at birth (P0) and found that 54.1% of the expected KO mice were present (72 KO / 133 WT), indicating that a substantial fraction of *Cnot6* KO mice died during the perinatal period.

Collectively, these findings suggest that CNOT6 is essential for neonatal and early postnatal growth, whereas CNOT6L plays a more prominent role in adult metabolic regulation.

### CNOT6 is highly expressed in postnatal livers and alters hepatic FGF21 and IGF1/IGFBP1 signaling

Given the observed defects in body and organ growth in *Cnot6* KO pups, we next analyzed the tissue-specific expression pattern of *Cnot6* mRNA in 2-week-old mice. Quantitative PCR revealed that *Cnot6* mRNA was highest in the liver followed by the kidney, brain, lung, and spleen (Figure 4A). Furthermore, publicly available RNA-seq data from the Mouse ENCODE transcriptome project (NCBI Gene ID: 104625) showed that hepatic *Cnot6* mRNA expression declines from fetal stage (E14.5) to adulthood (Figure 4B), suggesting a developmentally regulated role for CNOT6, particularly in the liver during early life.

**Figure 4.**
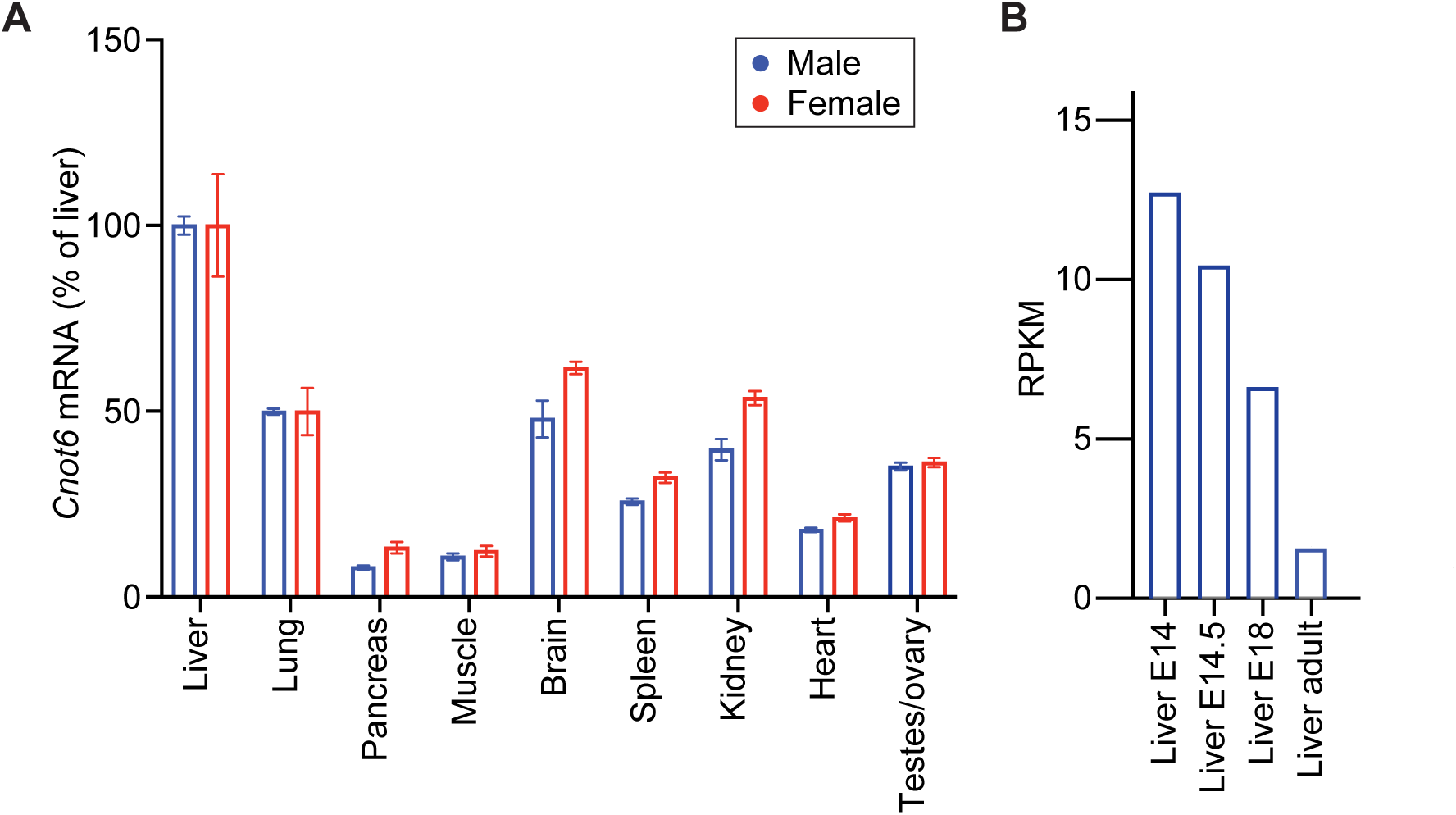
*Cnot6* mRNA is highly expressed in the liver during the perinatal and postnatal periods. (A) *Cnot6* mRNA levels in liver, lung, pancreas, muscle, brain, spleen, kidney, heart, and gonads (testes/ovary) of wild-type male and female mice at 2 weeks of age. Expression levels were normalized to *36B4* mRNA. n = 5-10 per group. (B) Developmental expression profile of hepatic *Cnot6* mRNA from embryonic day (E)14 to adulthood, obtained from publicly available RNA-seq data from the Mouse ENCODE transcriptome project. All data are presented as mean ± SEM.

**Figure 5.**
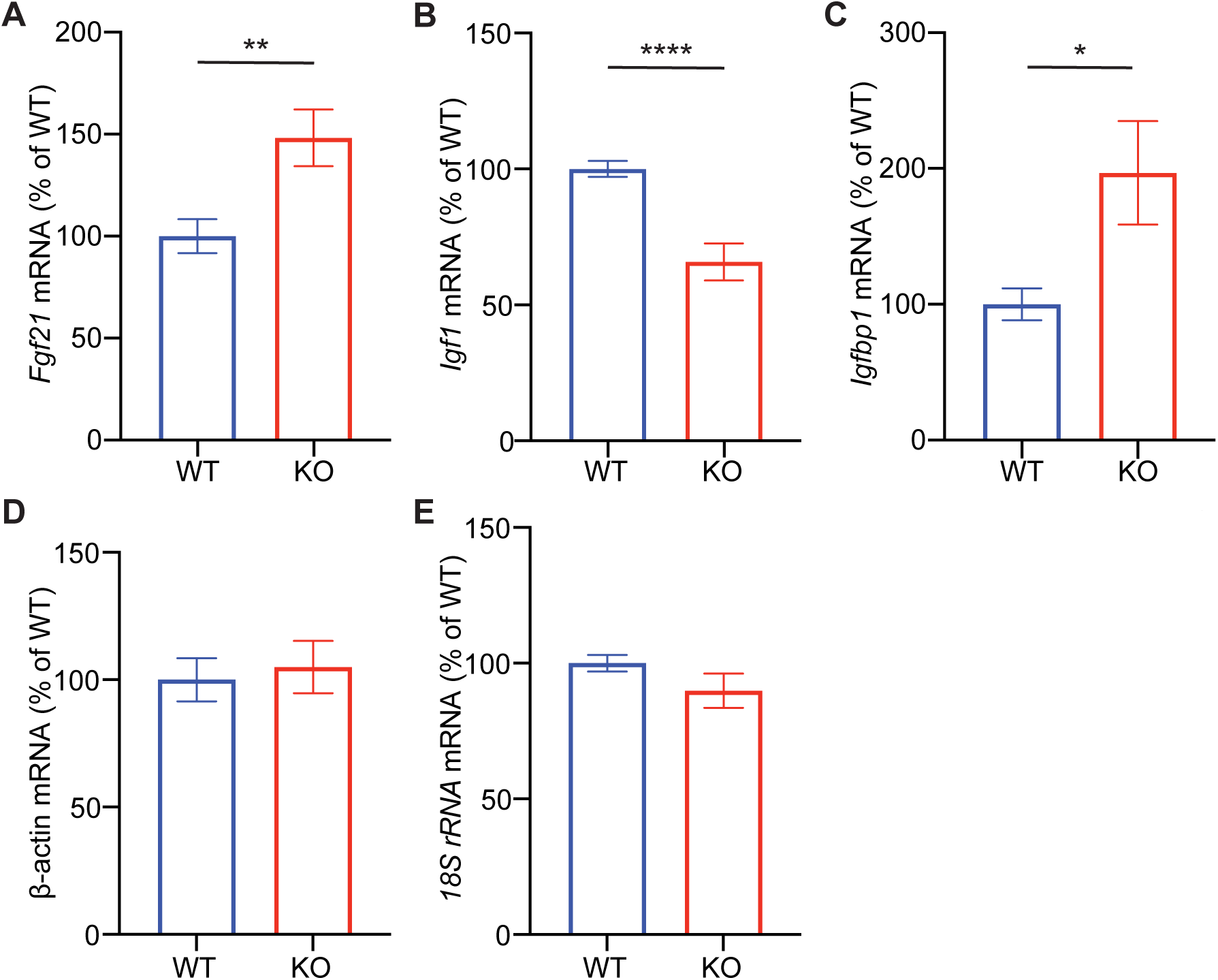
Increased *Fgf21* and *Igfbp1* expression and decreased *Igf1* expression in *Cnot6* KO livers. (A-C) Hepatic mRNA levels of *Fgf21* (A), *Igf1* (B), and *Igfbp1* (C) in male *Cnot6* WT and KO mice. (D-E) Expression of housekeeping genes *β-actin* (D) and *18S rRNA* (E) used as controls. All mRNA levels were normalized to 36B4 mRNA. n = 9-10 per group. All data are presented as mean ± SEM. *p < 0.05, **p < 0.01, ***p < 0.001, ****p < 0.0001. p values by the Student’s t test.

Since FGF21 is a known downstream target of CNOT6L in the adult liver (Katsumura et al., 2022; Morita et al., 2019), and has been implicated in growth suppression (Inagaki et al., 2008), we next investigated whether CNOT6 similarly regulates FGF21 in the postnatal liver. *Cnot6* KO mice exhibited significantly elevated hepatic *Fgf21* mRNA levels compared to controls (Figure 4D). Given that FGF21 suppresses IGF1 signaling during postnatal development (Inagaki et al., 2008), we examined components of this pathway. Consistent with elevated *Fgf21*, *Cnot6* KO livers showed significantly reduced *Igf1* mRNA and increased *Igfbp1* expression (Figures 4E–F), indicative of suppressed IGF1 signaling. Together, these findings suggest that CNOT6 regulates hepatic FGF21 expression and downstream IGF1-IGFBP1 signaling during postnatal development, thereby contributing to early-life growth control.

### Loss of *Cnot6* alters the hepatic transcriptome affecting metabolic and apoptotic programs

To investigate the molecular basis of the growth and metabolic abnormalities observed in *Cnot6*-deficient mice, we performed bulk RNA sequencing on liver tissues from 2-week-old *Cnot6* WT and KO mice. Principal component analysis (PCA) revealed distinct global transcriptomic profiles between WT and KO livers (Figure S4), indicating a substantial impact of *Cnot6* deletion on hepatic gene expression. Differential gene expression analysis (|log_2_FC| ≥ 1, adjusted p < 0.05) identified 378 differentially expressed genes (DEGs), including 203 upregulated and 175 downregulated genes (Figures 6A-B, Table S1). Consistent with prior qPCR results, key regulators, such as *Fgf21* and *Igfbp1*, were among the significantly altered genes (Figure 6B). Gene Set Enrichment Analysis (GSEA) revealed that downregulated genes were significantly enriched in pathways related to lipid biosynthetic processes and carbohydrate metabolism (Figure 6C). Conversely, upregulated genes were associated with apoptosis, cell proliferation, and stress responses (Figure 6D). Heatmap visualization highlighted altered expression of genes related to lipid metabolism (*Lipg*, *Cyp17a1*, *Mup3*), glucose metabolism (*Gck*, *Gpt*), and developmental regulation (Afp) (Figures 6E-G). In addition, stress- and apoptosis-related genes such as *Nr4a1*, *Pdcd5*, *Gzma*, and *Jun* were markedly elevated in KO livers (Figure 6H). These transcriptomic changes suggest that CNOT6 modulates metabolic and proliferation pathways in the liver, potentially through regulation of the FGF21-IGF1 signaling axis, contributing to the observed impairments in postnatal growth and metabolic homeostasis.

**Figure 6.**
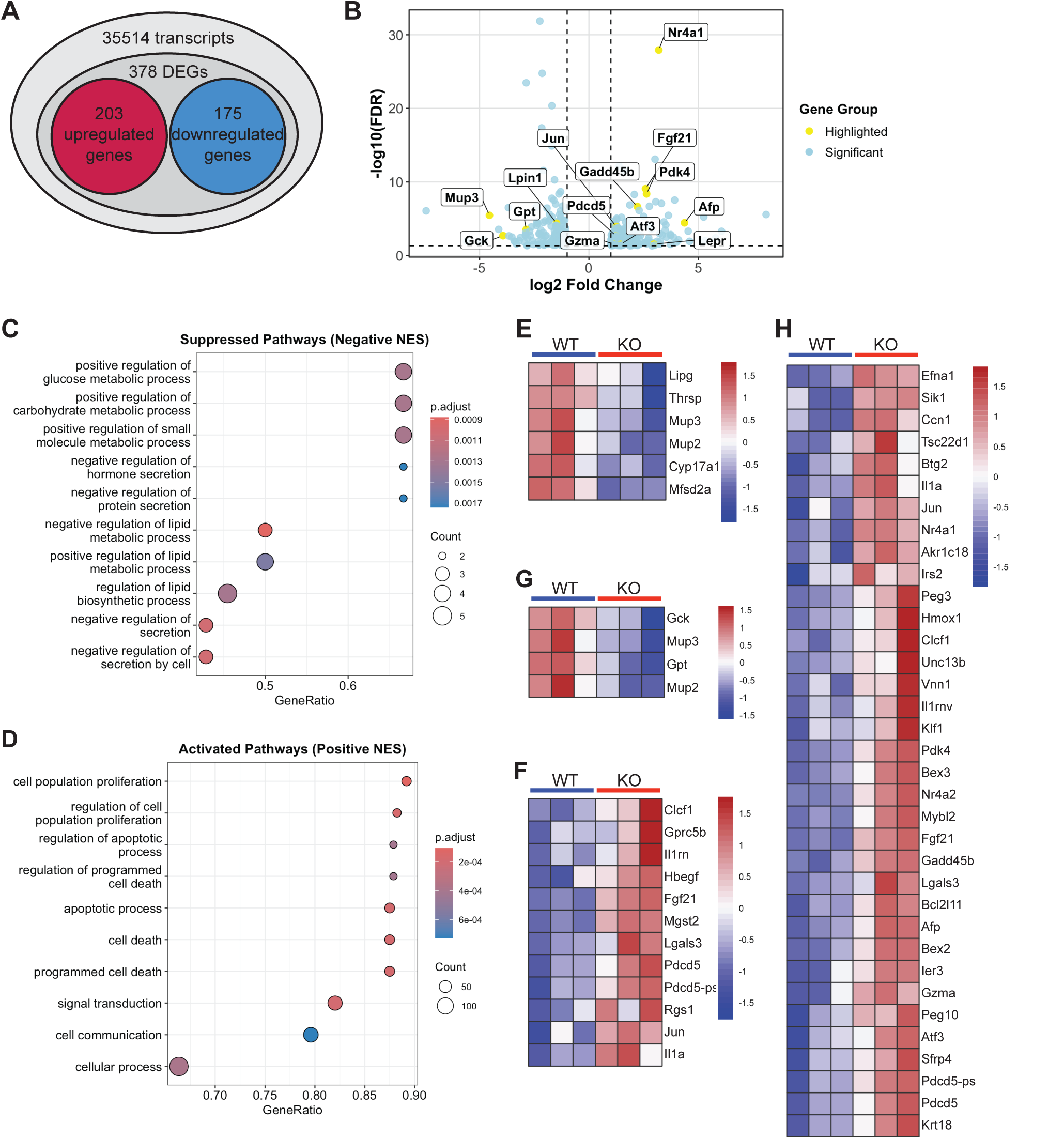
*Cnot6* deletion alters hepatic gene expression related to lipid and glucose metabolism and apoptosis. (A) Summary of RNA-seq results showing the number of upregulated and downregulated differentially expressed genes (DEGs) in *Cnot6* WT and KO livers. (B) Volcano plot displaying DEGs between *Cnot6* WT and KO livers. Blue dots represent significant DEGs, and selected genes are labeled in yellow. The x-axis indicates log_2_ fold change, and the y-axis shows - log_10_(FDR). Dashed vertical and horizontal lines indicate thresholds of |log_2_ fold change| ≥ 1 and adjusted p-value (FDR) ≤ 0.05, respectively. Volcano plots were generated using the ggplot2 package. (C-D) Dot plots showing the top ten significantly suppressed (C) and activated (D) pathways identified by Gene Set Enrichment Analysis (GSEA). GSEA results using clusterProfiler. (E-H) Heatmaps of representative DEGs from the identified pathways, including lipid biosynthetic process (E), apoptotic process (F), glucose metabolic process (G), and molecular function activator activity (H). Heatmaps were generated using the pheatmap package in R.

### Loss of *Cnot6* disrupts metabolic homeostasis during the postnatal stage

Building on transcriptomic evidence implicating altered lipid and glucose metabolism, we examined the metabolic impact of *Cnot6* deletion in postnatal mice. qPCR validation of RNA-seq-identified genes revealed consistent expression changes in pathways related to lipid and carbohydrate metabolism and cellular stress. In the liver of *Cnot6* KO mice, *Pdk4* mRNA, which diverts glucose away from oxidation and promotes fatty acid utilization, was markedly upregulated (Figure 7A). Similarly, expression of *Nr4a1* mRNA, a nuclear receptor induced by metabolic and inflammatory stress, and Gzma, a granzyme associated with apoptosis, was elevated. Conversely, several metabolic genes were significantly downregulated. These included *Lpin1*, a phosphatidic acid phosphatase critical for triglyceride synthesis and lipid storage; *Gck*, a key enzyme initiating hepatic glucose metabolism; *Gpt*, which links amino acid catabolism to gluconeogenesis; and *Mup3*, a member of the major urinary protein family that contributes to systemic energy regulation. To assess whether these transcriptional changes translated into physiological outcomes, we measured circulating β-hydroxybutyrate and blood glucose levels in 2-week-old animals. Consistent with a shift toward catabolic metabolism, *Cnot6* KO mice displayed elevated serum ketone bodies, indicating increased hepatic fatty acid oxidation. Interestingly, blood glucose levels were also elevated, potentially reflecting impaired glucose clearance or dysregulated insulin signaling.

**Figure 7.**
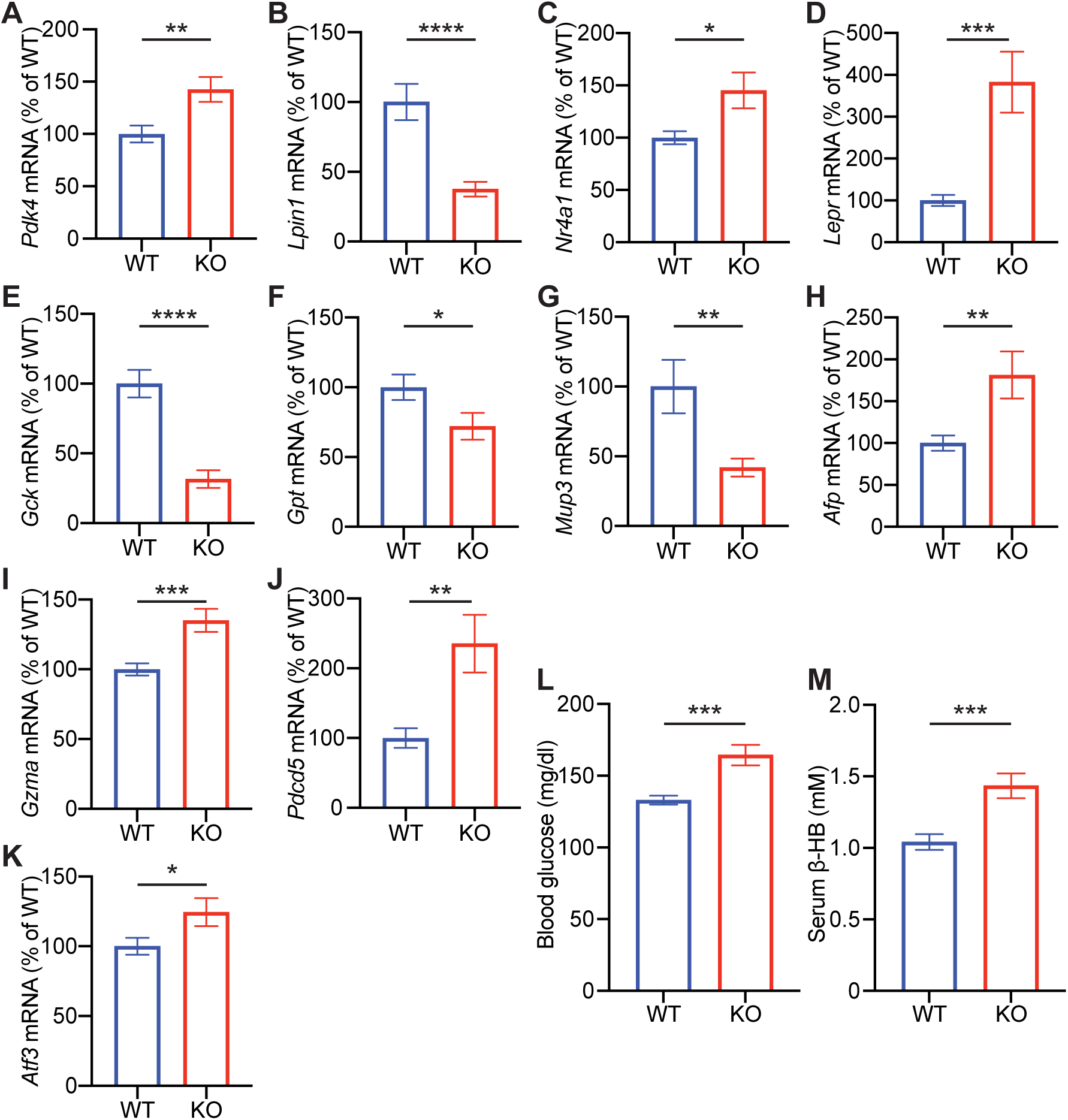
*Cnot6* deficiency drives a fasting-like, immature, and stress-responsive state in the liver. (A-M) Hepatic mRNA levels of *Pdk4* (A); *Lpin1* (B); *Nr4a1* (C); *Lepr* (D); *Gck* (E); *Gpt* (F); *Mup3* (G); *Afp* (H); *Gzma* (I); *Pdcd5* (J); and *Atf3* (K) in *Cnot6* WT and KO mice. (L-M) Levels of blood glucose (L) and serum β-hydroxybutyrate (ketone body) (M) in *Cnot6* WT and KO mice. n = 15-20 per group. All data are presented as mean ± SEM. *p < 0.05, **p < 0.01, ***p < 0.001, ****p < 0.0001. p values by the Student’s t test.

Taken together, these findings indicate that loss of *Cnot6* disrupts neonatal metabolic homeostasis by altering gene networks governing lipid oxidation, glucose handling, and stress responses. This reprogramming likely contributes to the growth suppression observed in *Cnot6*-deficient mice and indicates a role for the CNOT6-FGF21 axis in coordinating metabolism and development during the postnatal stage.

## DISCUSSION

This study identifies CNOT6 deadenylase, but not its paralog CNOT6L, as a key post-transcriptional regulator of early postnatal development acting through hepatic FGF21 expression. The CCR4-NOT complex, the central mRNA deadenylase machinery that determines transcript stability and turnover, has been primarily linked to adult metabolic control, yet its developmental functions have remained obscure. We demonstrate that *Cnot6* deficiency leads to elevated FGF21 and IGFBP1 levels, reduced IGF1 expression, and impaired postnatal growth, accompanied by increased ketone body production and enhanced expression of fatty acid oxidation-related genes. FGF21, previously characterized as a fasting-induced hepatokine that promotes energy expenditure in adults, here emerges as a mediator of neonatal growth regulation. These findings support a mechanistic link between mRNA decay and hormonal–metabolic pathways that coordinate anabolic and catabolic balance during the neonatal period. While the CNOT6L-FGF21 axis was previously implicated in adult metabolic regulation, our results uncover a developmentally distinct CNOT6-FGF21 pathway that governs growth and metabolic adaptation in early life, revealing a stage-specific specialization of the CCR4-NOT complex.

The CCR4-NOT complex acts as the major cytoplasmic deadenylase controlling the stability of metabolic transcripts. In the neonatal liver, our data suggest that CNOT6 regulates Fgf21 mRNA turnover, consistent with our previous finding that CNOT6L promotes Fgf21 decay in adult hepatocytes (Morita et al., 2019a; Katsumura et al., 2022a). Unlike CNOT6L, which is mainly expressed in adults, CNOT6 expression peaks during the early postnatal period, coinciding with the metabolic transition from placental to independent energy supply. Loss of CNOT6 leads to Fgf21 accumulation and suppression of the IGF1-IGFBP1 axis, a hormonal signature associated with growth inhibition. Although direct genetic evidence linking FGF21 to the *Cnot6* KO phenotype remains to be established, the post-transcriptional and metabolic changes observed in *Cnot6*-deficient mice are consistent with those previously described in *Cnot6l*-deficient mice, where FGF21-dependent mechanisms have been demonstrated (Katsumura et al., 2022a). Moreover, Fgf21 mRNA is directly targeted by the RNA-binding protein TTP, which recruits it to the CCR4-NOT complex in response to nutrient signals (Morita et al., 2019a). These findings suggest that CNOT6 likely regulates neonatal growth and metabolism through an FGF21-mediated pathway.

Although CNOT6 and CNOT6L share high sequence identity (>80%) and comparable enzymatic activity, our findings indicate that they are functionally segregated in a developmental stage-dependent manner. These paralogs are mutually exclusive within the CCR4-NOT complex, such that each complex incorporates either CNOT6 or CNOT6L as its catalytic subunit, but not both. During neonatal growth, CNOT6 is predominantly expressed and supports anabolic metabolism required for rapid tissue expansion, whereas CNOT6L becomes more active in adulthood, coordinating lipid oxidation and energy expenditure to maintain systemic homeostasis. This isoform transition within the CCR4-NOT complex likely reprograms the mRNA decay landscape to align with the shifting metabolic priorities between developmental stages—promoting biosynthetic and growth programs early in life and enforcing metabolic homeostasis later on. Such stage-specific utilization of distinct deadenylase subunits may represent an adaptive mechanism that enables the CCR4-NOT complex to dynamically tune post-transcriptional regulation according to physiological demands.

The high expression of *Cnot6* mRNA in fetal and neonatal liver, followed by its decline in adulthood, indicates a transient yet essential role in orchestrating the metabolic transition from placental to autonomous energy supply. This developmental window represents a critical phase in which lipid and carbohydrate fluxes, together with endocrine signals such as GH and IGF1, must be precisely coordinated to maintain the balance between anabolic and catabolic metabolism required for rapid tissue expansion. Our findings suggest that CNOT6-mediated repression of FGF21 acts as a molecular safeguard that prevents premature activation of catabolic programs, thereby preserving the anabolic environment necessary for growth. The severe growth retardation and early postnatal lethality observed in *Cnot6*-deficient mice underscore the physiological importance of this control mechanism. From an evolutionary perspective, this regulatory axis may have emerged as a conserved checkpoint ensuring survival during the critical transition when nutrient supply shifts from maternal to external sources. By coupling post-transcriptional regulation to hormonal and metabolic reprogramming, the CNOT6-FGF21 axis provides a flexible mechanism to balance growth, energy conservation, and stress resilience during early postnatal life.

Clinically, dysregulation of the FGF21 axis has been linked to impaired growth and metabolic imbalance in humans. Elevated circulating FGF21 levels are observed in preterm infants with postnatal growth failure (Guasti et al., 2014; Mistry et al., 2023), in children with idiopathic short stature or growth hormone deficiency (Lee et al., 2023), and in adolescents with anorexia nervosa (Fazeli et al., 2010). All of these conditions are characterized by GH resistance and reduced IGF1 signaling. These clinical patterns mirror several aspects of our findings in *Cnot6*-deficient mice, in which excessive FGF21 suppresses IGF1 signaling while increasing IGFBP1 and blunts postnatal growth. Notably, nutritional supplementation alone often fails to restore normal growth in these patients, which highlights the need to identify additional molecular mechanisms that coordinate growth and metabolism. Together, these observations suggest that CNOT6-mediated regulation of FGF21 represents a conserved mechanism that maintains anabolic balance during development. Importantly, targeting this pathway, either by restoring CNOT6 activity or by fine-tuning CCR4-NOT dependent mRNA decay, could provide new therapeutic opportunities for growth failure and metabolic disorders that remain refractory to current nutritional or GH-based interventions.

## Materials and Methods

### Generation and maintenance of *Cnot6* KO mice

A *Cnot6* KO mouse line (*Cnot6^tm1a(KOMP)Wtsi^*) (Figure S2) was obtained as frozen sperm from the Mutant Mouse Resource and Research Centers (MMRRC) supported by the NIH. The frozen sperm were used for *in vitro* fertilization (IVF) with oocytes from C57BL/6N females to generate founder mice. The line was subsequently backcrossed to the C57BL/6J background for more than 10 generations. Mice were maintained under a 12-hour light-dark cycle with free access to food and water. Genotypes were determined by PCR using specific primers. Body weight was recorded at birth (postnatal day 0) and weekly thereafter until study completion. For tissue collection, mice were euthanized at 2 weeks of age under isoflurane anesthesia, followed by terminal cardiac puncture. Tissues, including liver, lung, heart, pancreas, spleen, kidney, stomach, gastrocnemius muscle, tibia, and brain, were collected and immediately flash-frozen in liquid nitrogen. Blood glucose levels were measured immediately before euthanasia using a Contour Next test strip (Ascensia Diabetes Care AG). Serum was collected by centrifugation and stored at -80°C. All experiments were conducted in accordance with the guidelines for animal use and approved by the Institutional Animal Care and Use Committee (IACUC) of the University of Texas Health Science Center at San Antonio.

### Serum Assays

Mouse blood was collected at the end of the study via terminal cardiac puncture. Whole blood was allowed to clot at room temperature and centrifuged at ∼1,000×g for 30 min at 4°C to obtain serum. Serum samples were stored at -80°C and thawed on ice immediately before analysis. Serum β-hydroxybutyrate (ketone bodies) concentrations were measured using a β-Hydroxybutyrate Colorimetric Assay Kit (Cayman Chemical) according to the manufacturer’s instructions. Absorbance was read using a Cytation 1 plate reader (Agilent Technologies).

### Western Blot

Mouse tissues were homogenized in lysis buffer (50 mM Tris–HCl, pH 7.5, 150 mM NaCl, 1 mM EDTA, 1% NP-40, and protease inhibitor cocktail [Roche]) using a glass homogenizer. Lysates were centrifuged at 12,000×g for 10 min at 4°C, and supernatants were collected. Protein concentrations were determined using the BCA Protein Assay Kit (Thermo Fisher Scientific) and quantified with a Cytation 1 plate reader (Agilent Technologies). Equal amounts of protein (50 µg per lane) were resolved by SDS-PAGE and transferred onto PVDF membranes (Thermo Fisher Scientific). Membranes were blocked in 5% skim milk in TBST and incubated overnight at 4°C with primary antibodies against CNOT6 (home-made) and β-actin (Cell Signaling Technology, #4970). After washing, membranes were incubated with HRP-conjugated anti-rabbit IgG (Jackson ImmunoResearch) and developed using enhanced chemiluminescence (ECL, Thermo Fisher Scientific).

### RNA isolation and RT-qPCR

Total RNA was isolated from livers of 2-week-old *Cnot6* WT and KO mice using TRIzol reagent (Thermo Fisher Scientific) according to the manufacturer’s protocol. RNA concentration and purity were determined using a NanoDrop One^C^ spectrophotometer (Thermo Fisher Scientific). 2 μg of total RNA was reverse transcribed using the High-Capacity cDNA Reverse Transcription Kit (Applied Biosystems). Quantitative PCR (qPCR) was performed on a StepOnePlus Real-Time PCR System (Applied Biosystems) in 96-well format using PowerUp SYBR Green Master Mix (Applied Biosystems). Relative mRNA expression levels were calculated using standard curves generated from serially diluted cDNA templates, with murine *ribosomal protein Rplp0* (*36B4*) as the internal reference gene. Primer sequences are listed in Table S2.

### Bulk RNA-seq Library Preparation and Sequencing

Total RNA was extracted from livers of 2-week-old *Cnot6* WT and KO mice using TRIzol reagent (Thermo Fisher Scientific). RNA quantity and quality were assessed using a NanoDrop One^C^ spectrophotometer and Agilent TapeStation, with all samples exhibiting RNA Integrity Numbers (RIN) greater than 9.0. Libraries were prepared from three independent biological replicates per genotype using the TruSeq Stranded mRNA LT Sample Prep Kit (Illumina) following the manufacturer’s instructions. Sequencing was performed on an Illumina NovaSeq 6000 platform to generate paired-end reads at the Sequencing Facility at University of Texas Health Science Center at San Antonio.

### RNA-seq data processing and analysis

Raw sequencing reads were first assessed for quality using FastQC (v0.11.9). Adapter sequences and low-quality bases were trimmed using Trimmomatic (v0.39) with the parameters: ILLUMINACLIP:2:30:10, LEADING:3, TRAILING:3, SLIDINGWINDOW:4:15, and MINLEN:36. Filtered reads were aligned to the Mus musculus reference genome (GRCm39, Ensembl release 104) using HISAT2 (v2.2.1) with the --dta option. Alignment files were converted and sorted into BAM format using SAMtools (v1.16.1). Transcript assembly and quantification were performed using StringTie (v2.2.1) with default settings. The analysis pipeline was applied to three independent *Cnot6* WT and KO biological replicates.

Subsequent data analysis was conducted in R. Gene-level read counts were generated using featureCounts (Subread package). Genes with total counts ≤1 across all samples were excluded to remove low-abundance transcripts lacking statistical power for differential expression testing. Differential expression analysis was performed with DESeq2 (v1.49.3) (Love et al., 2014) using a model design of ∼ Genotype. Genes with an absolute log₂ fold change ≥1 and an adjusted p-value ≤0.05 were considered significantly differentially expressed. Regularized log transformation was applied for visualization. Functional enrichment analysis of differentially expressed genes was conducted using clusterProfiler (v3.10.1) (Yu et al., 2012) with Gene Ontology (GO) biological process annotations. Gene count matrices (Table S3) and the complete DEG list (Table S1) are provided. RNA-seq data have been deposited in the NCBI Gene Expression Omnibus (GEO).

### Quantification and statistical analysis

All quantitative data are presented as mean ± SEM. Statistical analyses were performed using GraphPad Prism 10. Comparisons among multiple groups were conducted using one-way ANOVA followed by Dunnett’s post hoc test or two-way ANOVA followed by Bonferroni’s post hoc test, as appropriate. For comparisons between two groups, unpaired two-tailed Student’s t-tests were used. Differences were considered statistically significant when *p* < 0.05.

## ACKNOWLEDGMENTS

We thank Luiz Penalva for assistance with RNA-seq analysis; Michelle Ramirez for help with mouse experiments; and Sherry Dodds for support. We also acknowledge the Animal Facility and Genome Sequencing Facility at UTHSCSA for RNA-seq and technical assistance. This work was supported by the Cancer Prevention and Research Institute of Texas (CPRIT) Awards (RP220267 and RP250118); NIH grant R01 AA031407; the American Cancer Society Research Scholar Grant (RSG-22-016-01-CCB); the Max and Minnie Tomerlin Voelcker Fund Young Investigator Award; the University of Texas Rising STARs Award; the Helen F. Kerr Foundation Grant; the Shelby Tengg Foundation Grant; the Cancer Center Support Grant (NIH P30 CA054174); the Grant-in-Aid for Scientific Research (25K02207); and the JST FOREST Program (JPMJFR216D). S.K. was supported by the Pepper Center RL5 Research Career Development Award (P30 AG044271), the American Heart Association Postdoctoral Fellowship, the JSPS Overseas Research Fellowship, and the Uehara Memorial Foundation Postdoctoral Fellowship.

## AUTHOR CONTRIBUTIONS

M.S., S.K., and M.M. conceived and designed the study and performed the experiments. M.S., S.K., and M.M. wrote the manuscript with input from all authors. M.Z. and D.G. assisted with experiments. All authors analyzed the data, discussed the results, and approved the final version of the manuscript.

## Supplemental Information

**Figure S1.**
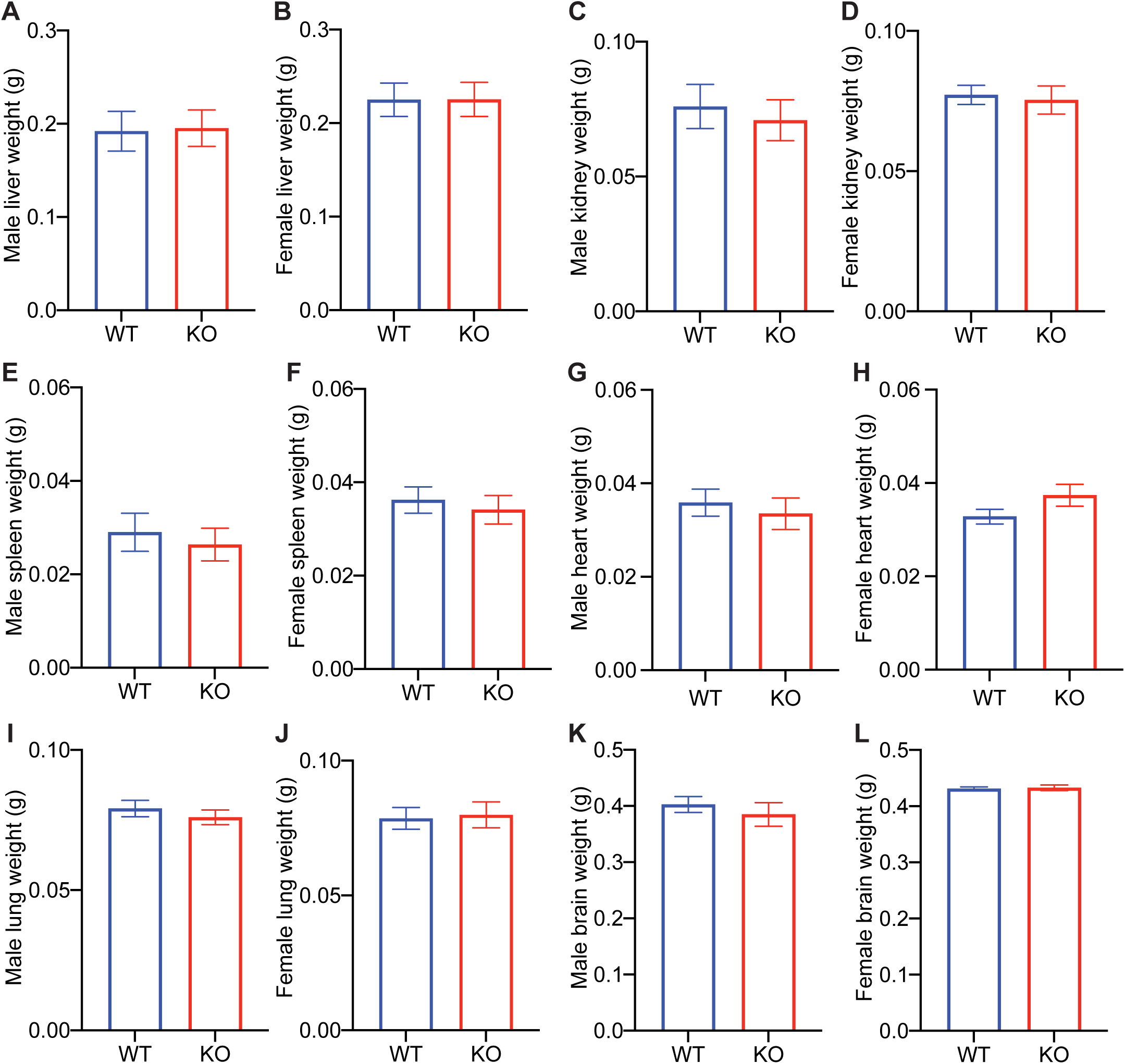
*Cnot6l* deletion does not affect organ growth during postnatal development. (A-L) Weights and representative images of tissues, including livers (A-B), kidneys (C-D), spleens (E-F), hearts (G-H), lungs (I-J), and brains (K-L) of *Cnot6l* WT and KO male and female mice. n = 5-10 per group. All data are presented as mean ± SEM. Related to Figure 1.

**Figure S2.**
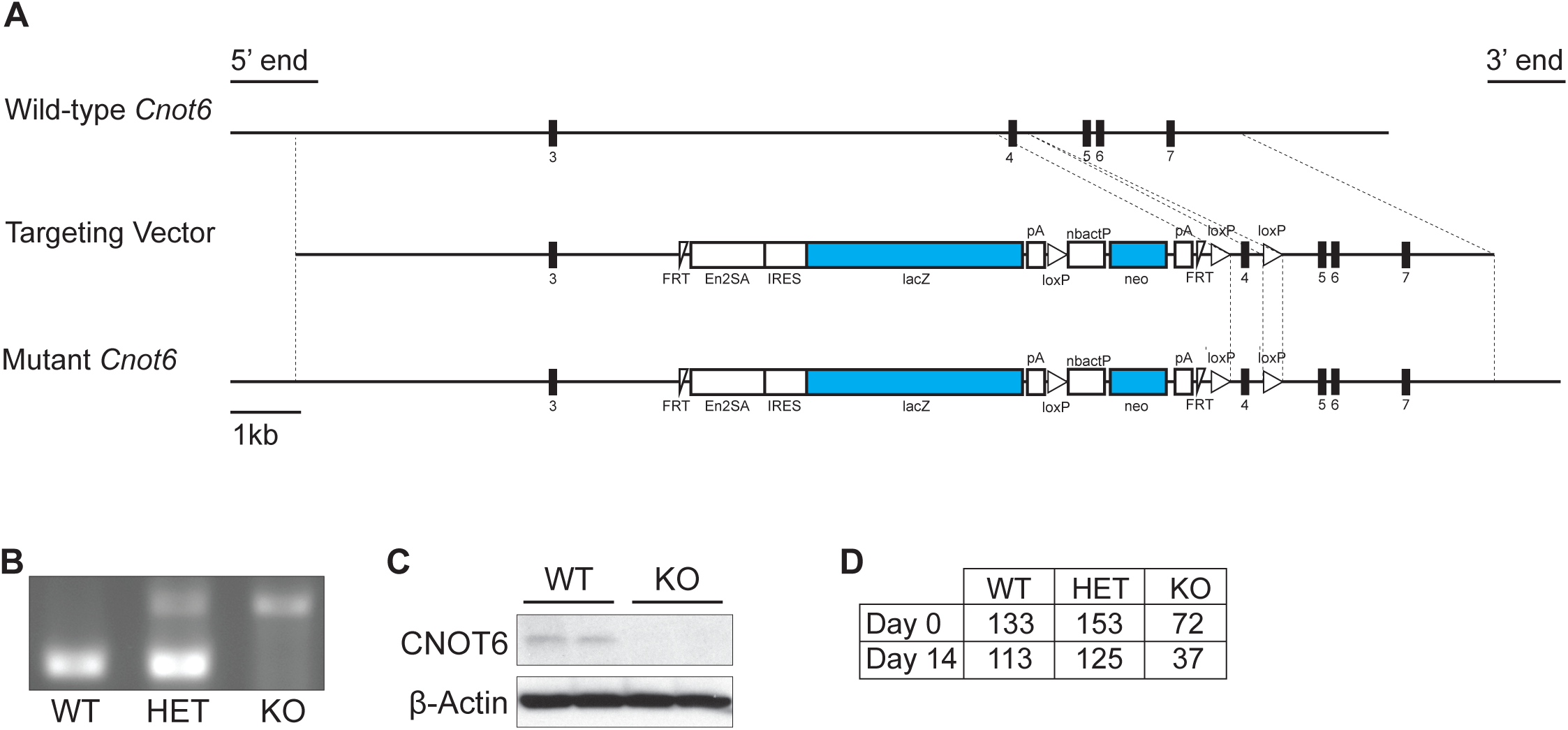
Generation and validation of *Cnot6* KO mice and increased perinatal lethality. (A) Schematic representation of the *Cnot6* WT allele, targeting vector, and mutant allele. White triangles indicate loxP sites. Blue boxes represent reporter and selection cassettes (lacZ, β-galactosidase; neo, neomycin). Numbered black boxes indicate *Cnot6* exons. The targeting vector was designed to disrupt normal transcription through a gene-trap strategy, resulting in complete loss of detectable CNOT6 protein. (B) Genotyping PCR results for *Cnot6* WT, HET, and KO mice. (C) Western blot analysis confirming loss of CNOT6 protein expression in *Cnot6* KO mice. β-actin was used as a loading control. (D) Genotype distribution of live pups from *Cnot6* heterozygous intercrosses at day 0 (birth) and day 14, showing reduced survival of homozygous KO mice. Related to Figure 2.

**Figure S3.**
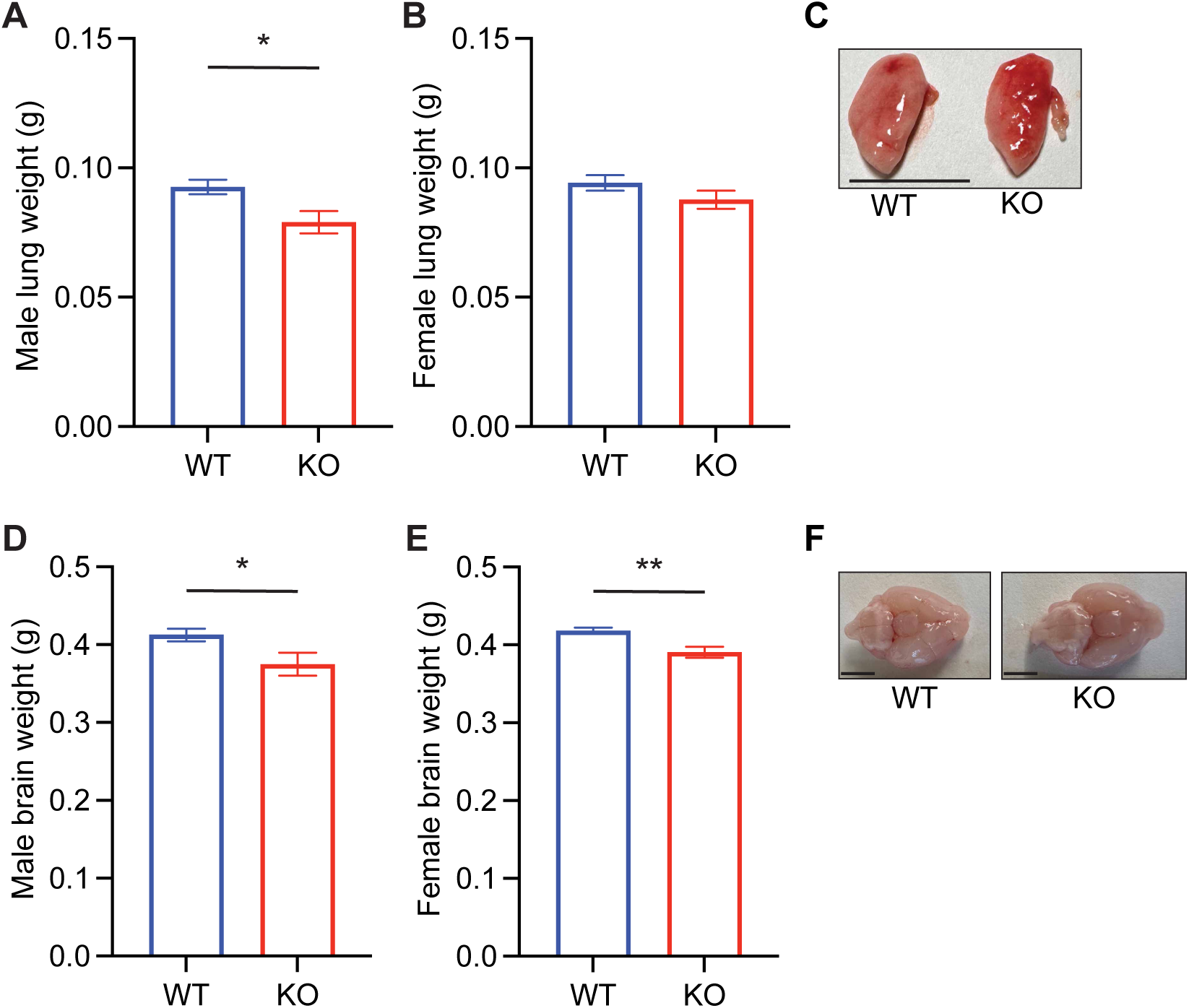
*Cnot6* deletion impairs organ growth during postnatal development. (A-F) Weights and representative images of tissues, including lungs (A-C) and brains (D-F) of *Cnot6* WT and KO male and female mice. n = 5-38 per group. Red reference line indicates 10 mm. All data are presented as mean ± SEM. *p < 0.05, **p < 0.01, ***p < 0.001, ****p < 0.0001; p values by the Student’s t test. Related to Figure 3.

**Figure S4.**
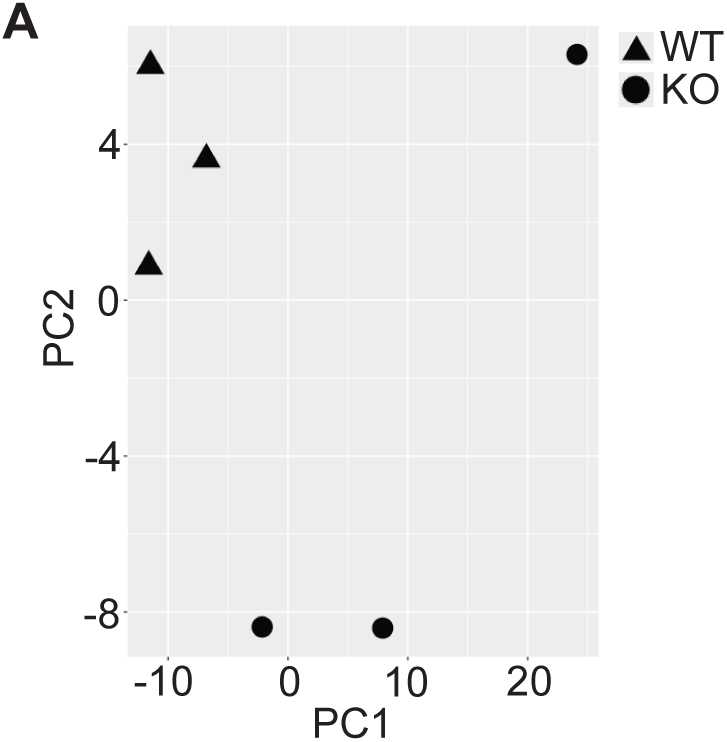
RNA-seq analysis of *Cnot6* WT and KO livers. (A) Principal Component Analysis (PCA) of male Cnot6 WT and KO liver RNA-seq datasets. Each point represents one biological replicate. PCA was performed using the plotPCA function in R based on log_2_-transformed normalized gene counts. WT and KO samples form distinct clusters, indicating clear genotype-dependent transcriptional profiles.

**Table S1. Differentially expressed genes in livers of 2-week-old *Cnot6* WT and KO mice**

Differential expression analysis was performed using DESeq2. Genes with |log₂ fold change| ≥ 1 and adjusted p ≤ 0.05 were considered significant (n = 378).

**Table S2. Primer sequences used for qPCR**

Forward and reverse primers used in this study.

**Table S3. Gene-level RNA-seq read counts from Cnot6 WT and KO mouse livers**

Gene-level raw read counts obtained from bulk RNA-seq analysis of liver tissues from 2-week-old *Cnot6* WT (WT1-WT3) and KO (KO1-KO3) mice. Counts represent unnormalized read numbers prior to differential expression analysis.

